# Comparison of transparency and shrinkage of insect brains using novel and traditional clearing methods

**DOI:** 10.1101/2020.08.28.273128

**Authors:** Bo M. B. Bekkouche, Helena K. M. Fritz, Elisa Rigosi, David C. O’Carroll

**Affiliations:** Department of Biology, Lund University, Lund, Sweden

**Author notes:** Correspondence: Bo B. M. Bekkouche, Department of Biology, Lund University, Sölvegatan 35, 22362 Lund, Sweden.

**Keywords:** Optical tissue clearing, transparency, shrinkage, insect brain, brain imaging, neuron imaging, super-resolution microscopy, digital light-sheet

## Abstract

Improvement of imaging quality has the potential to visualize previously unseen building blocks of the brain and is therefore one of the great challenges in neuroscience. Rapid development of new tissue clearing techniques in recent years have attempted to solve imaging compromises in thick brain samples, particularly for high resolution optical microscopy, where the clearing medium needs to match the refractive index of the objective immersion medium. These problems are exacerbated in insect tissue, where numerous (initially air-filled) tracheal tubes branching throughout the brain increase the scattering of light. To date, surprisingly few studies have systematically quantified the benefits of such clearing methods using objective transparency and tissue shrinkage measurements. In this study we compare a traditional and widely used insect clearing medium, methyl salicylate combined with permanent mounting in Permount (‘MS/P’) with several more recently applied clearing media that offer tunable refractive index (*n*): 2,2-thiodiethanol (TDE), ‘SeeDB2’ (in variants SeeDB2S & SeeDB2G matched to oil and glycerol immersion, *n*=1.52 & 1.47 respectively) and Rapiclear (also with *n*=1.52 & 1.47). We measured transparency and tissue shrinkage by comparing freshly dissected brains with cleared brains from dipteran flies, with or without addition of vacuum or ethanol pre-treatments (dehydration and rehydration) to evacuate air from the tracheal system. The results show that ethanol pre-treatment is very effective for improving transparency, regardless of the subsequent clearing medium, while vacuum treatment offers little measurable benefit. Ethanol pre-treated SeeDB2G and Rapiclear brains show much less shrinkage than using the traditional MS/P method. Furthermore, these newly developed media offer outstanding transparency compared to TDE and MS/P. Rapiclear protocols were less laborious compared to SeeDB2, but both offer sufficient transparency and refractive index tunability to permit super-resolution imaging of local volumes in whole mount brains from large insects, and even light-sheet microscopy. Although long-term permanency of Rapiclear stored samples remains to be established, our samples still showed good preservation of fluorescence after storage for more than a year at room temperature.

## 1 Introduction

A number of technological advances in recent years have increased the desirability of imaging structures within the brain in high resolution using optical rather than traditional histological sectioning. Applications range from imaging neurons revealed via optogenetic techniques, to *in-situ* neuronal tracing in order to establish circuit level interactions between neurons (Dunbier et al., 2012; Keles and Frye, 2017). In our own lab, for example, we are interested in applying biophysically realistic computational models to improve our understanding and test hypotheses regarding how the brain works (Shoemaker, 2011; Bekkouche et al., 2017). Such models rely on high resolution microscopy, enabling 3D reconstruction of individual neuron morphologies within the brain, to establish the likely biophysical compartments through which neuronal signals travel. One general hypothesis in this type of modeling is that the fine neurites perform computations that are important for individual neuron function, which in turn is part of a neural network connecting to the rest of the brain. The higher the imaging quality, the greater the certainty that the morphology of all fine neurites and likely synaptic zones that contribute to the response are faithfully captured by the imaging system and can thus be incorporated into the model.

One challenge for the application of such *in-situ* optical imaging techniques, however, is the limitation to resolution caused by light scattering within the thick tissue sample. This requires the application of a suitable clearing medium (Silvestri et al., 2016; Ariel, 2017) to reduce local refractive index differences that provide the basis for such scattering (Schmitt and Kumar, 1998; Tuchin et al., 1998; Tuchin, 2005; Harvard et al., 2015). These problems are exacerbated in insect tissue by their tracheal respiratory system: numerous air-filled tracheal tubes branch finely throughout the brain increasing these index differences and thus scattering of light.

A second challenge for *in-situ* neuronal imaging is the tradeoff between the refractive index of the objective immersion medium and the working distance: The high numerical apertures required for high-resolution microscopy depend on a high refractive index immersion medium. However, this typically comes with a compromise of short working distance, often <100µm, making traditional oil immersion objectives (with the highest theoretical resolution) less compatible with imaging neurons within very thick tissue samples which require long working distance to focus deeper into the tissue. Such objectives are entirely unsuitable for whole-mount imaging of large organs such as rodent brains. Even in insects such as those studied in our own lab, the brains of larger species such as dipteran flies, bees and dragonflies are approximately 500µm thick even in their smallest dimension.

There are several possible solutions to achieving both the high numerical aperture required for high or super-resolution imaging and the large working distance required for imaging brains in whole mount. Several microscope manufacturers have now developed direct immersion objectives optimized for specific clearing media and with working distances up to several mm, permitting direct imaging of cleared brains without a coverslip (Hama et al., 2015; Liu et al., 2016). However, the very large size of such objectives and the specific match to a single immersion medium requires an upright microscope system dedicated to the goal of imaging this specific tissue type. Furthermore, without a coverslip, such samples may be less stable, either for super-resolution imaging *in situ*, or for long term storage. Water immersion objectives with very long working distance provide an alternative more suitable for application to general purpose microscopes, but lack the outright numerical aperture required for super-resolution imaging, and suffer from a mismatch to the refractive index of typical (non-aqueous) clearing media, leading to spherical aberration that increases with imaging depth (Dong et al., 2004; Gonzalez-Bellido and Wardill, 2012). As a compromise between these extremes, high numerical aperture objectives designed to work with cover-slipped samples have recently been produced by several vendors, using intermediate index immersion media such as silicone oil or glycerol. In some cases (e.g. the Leica HC PL APO CS2 63x/1.3 glycerol objective used in the present paper, free working distance=300µm) these come close to the highest numerical aperture achieved by oil-immersion objectives, but offer up to several times the effective working distance. One study recently demonstrated that by combining such objectives with a novel clearing medium matched to the refractive index of glycerol, it is possible to apply super-resolution techniques such as STED or deconvolution to fine neurites more than 100µm below the sample surface (Ke et al., 2016).

In insect neurobiology, several traditional clearing media such as terpiniol and methyl salicylate have been widely applied for whole-mount imaging of Golgi stained or fluorescently labelled samples (Kurylas et al., 2008; el Jundi et al., 2010; Lin and Strausfeld, 2012; Stöckl and Heinze, 2015; Adden et al., 2020). The dehydration of the brain with organic solvents required before clearing in such media generally aids displacement of air trapped in the tracheal system, leading to good tissue clarity. In many labs, it has become standard practice to permanently mount such methyl salicylate cleared samples using a hardening mounting medium such as Permount™, before imaging. This permits the very stable imaging required to combine multiple large volume image sets for whole brain reconstructions or for tracing neurons with processes that may travel up to several millimeters across the brain (Geurten et al., 2007; Kurylas et al., 2008; el Jundi et al., 2010). Another advantage of this method is that samples then survive storage for many years, and can subsequently be re-imaged using newly available techniques or even re-treated with new immunohistochemical analyses after dissolving out the mounting medium in a suitable solvent (Heinze and Homberg, 2008; Heinze et al., 2013).

While clearing in methyl salicylate/ Permount is widely used, this technique is not without problems for insect brain imaging (Gonzalez-Bellido and Wardill, 2012). In particular, the high refractive index of this medium (1.52) requires the use of oil immersion objectives in order to avoid spherical aberration within the sample. As described earlier, this limits the useful depth that can be imaged, and often requires samples to be flipped and re-imaged from both sides. This complicates combining the imaged volumes before subsequent analysis, even if using lower numerical aperture oil objectives. The limited working distance of oil-immersion objectives makes such media incompatible with super-resolution imaging of anything other than superficial structures unless tissue is subjected to traditional histological sectioning. As an alternative, Gonzalez-Bellido & Wardill (Gonzalez-Bellido and Wardill, 2012) used 2,2-thiodiethanol (TDE) to clear insect tissue, a medium that permits refractive index tuning via dilution with small volumes of water.

While TDE may provide a suitable solution in some applications and can provide an excellent optical point-spread function deep within an embedded sample, it is known to quench the fluorescence of many fluorophores (Ke et al., 2016). This has led to a number of efforts to develop alternative tunable media to match the refractive index of the required immersion medium (e.g. silicone oil or glycerol) for long working distance objectives. Ke et al. (2016) recently introduced a novel non-quenching clearing medium, ‘SeeDB2’, based on iohexol, in variants that can either match immersion oil or glycerol. They demonstrated an excellent 3-dimensional point spread function, rivalling that of TDE and then successfully applied this to several super-resolution imaging applications. Another new clearing medium recently introduced to insect brain research (Frank et al., 2017), Rapiclear (developed by Sunjin lab Co.), also has the advantage of tunable refractive index. Media such as Rapiclear and SeeDB2 are both water soluble, so do not require a final dehydration stage before mounting and imaging. This potentially gives better control over tissue shrinkage. Tissue shrinkage is, of course, highly undesirable for super-resolution imaging or if the goal is to obtain accurate estimates of neurite diameters, as required for accurate compartment modeling of neurons.

To date there has been no systematic analysis of the pros and cons and cross-comparison of these traditional or recently developed methods using objective measurements of transparency and shrinkage. Many clearing media comparison studies have lacked application of a mathematically based transparency measurement method (Liu et al., 2016; Yu et al., 2018). This could introduce subjectivity into the estimation of transparency. Loren et al. (2019) focused on comparing attenuation of transparency over time using an objective measurement of the transparency of different clearing media. Yu et al. (2018) discussed several new clearing media, but the transparency was not quantified between clearing techniques. Wan et al. (2018) compared seven clearing methods using a mathematical approach to calculate imaging depth based on intensity which could be considered as a measurement of transparency. However, none of these studies systematically quantified differences in shrinkage, and none of them focused specifically on insect tissue, with the additional challenges presented by the tracheal respiratory system.

In this paper we developed a standard experiment protocol to quantify the trade-off between tissue transparency, final refractive index and shrinkage or expansion artefacts using several of these recently developed clearing methods. We benchmarked these against the more traditional methyl salicylate / Permount method. We measured transparency of brain images against a background grating to estimate the contrast attenuation by the tissue, then estimated shrinkage by identifying common reference points in images of large fly brains (from hoverflies and blowflies) before and after clearing. We found that among the new clearing methods, both SeeDB2 and Rapiclear not only reduce shrinkage compared with methyl salicylate cleared brains, they also provide a substantially more transparent brain. In addition, the compatibility of Rapiclear with a range of direct-immersion objectives makes this an outstanding choice for new applications such as light-sheet microscopy.

## 2 Materials and Equipment

Table 1 provides a list of reagents/ materials and/or equipment specifically required for the methods we developed.

**Table 1.**
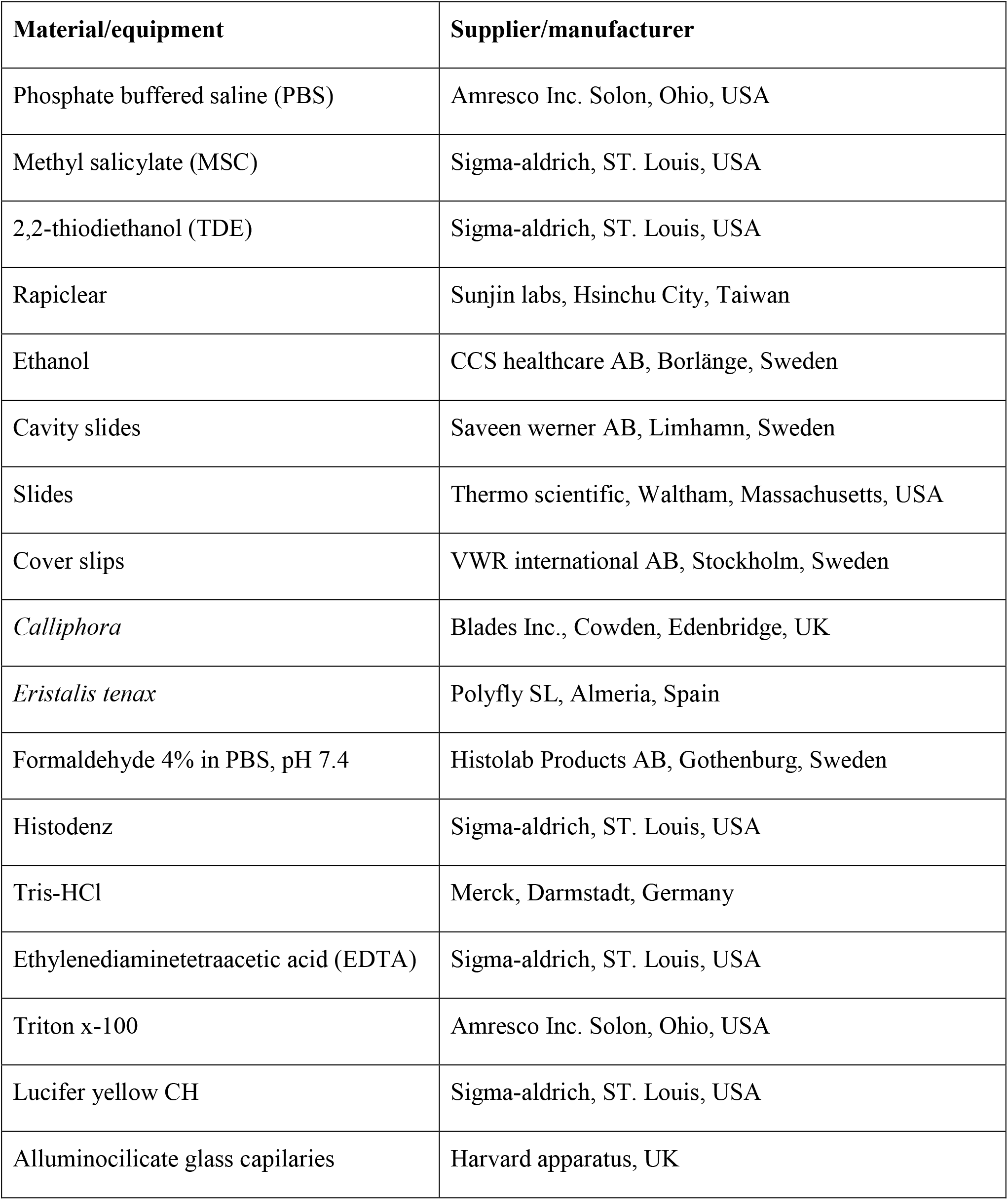
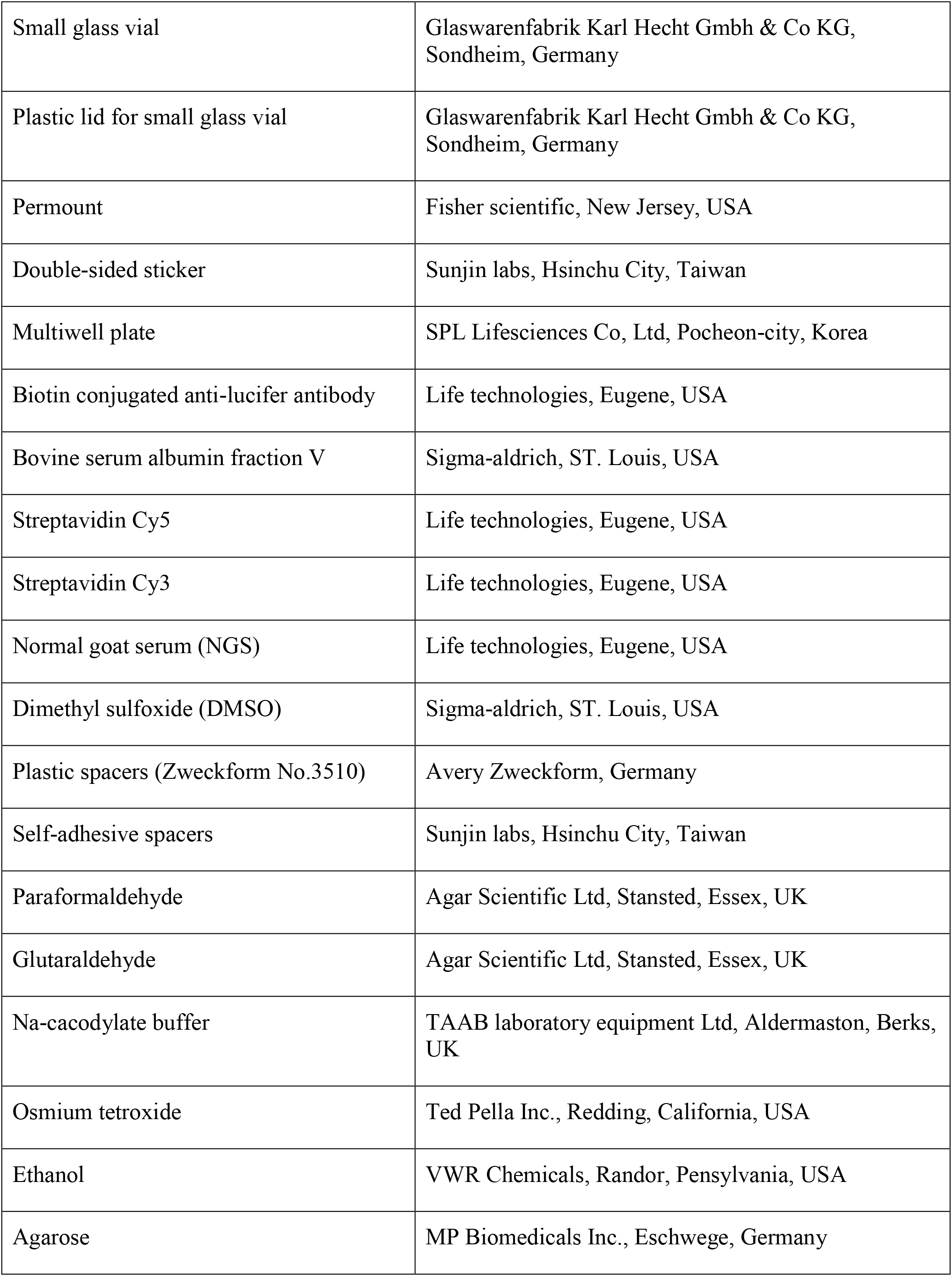
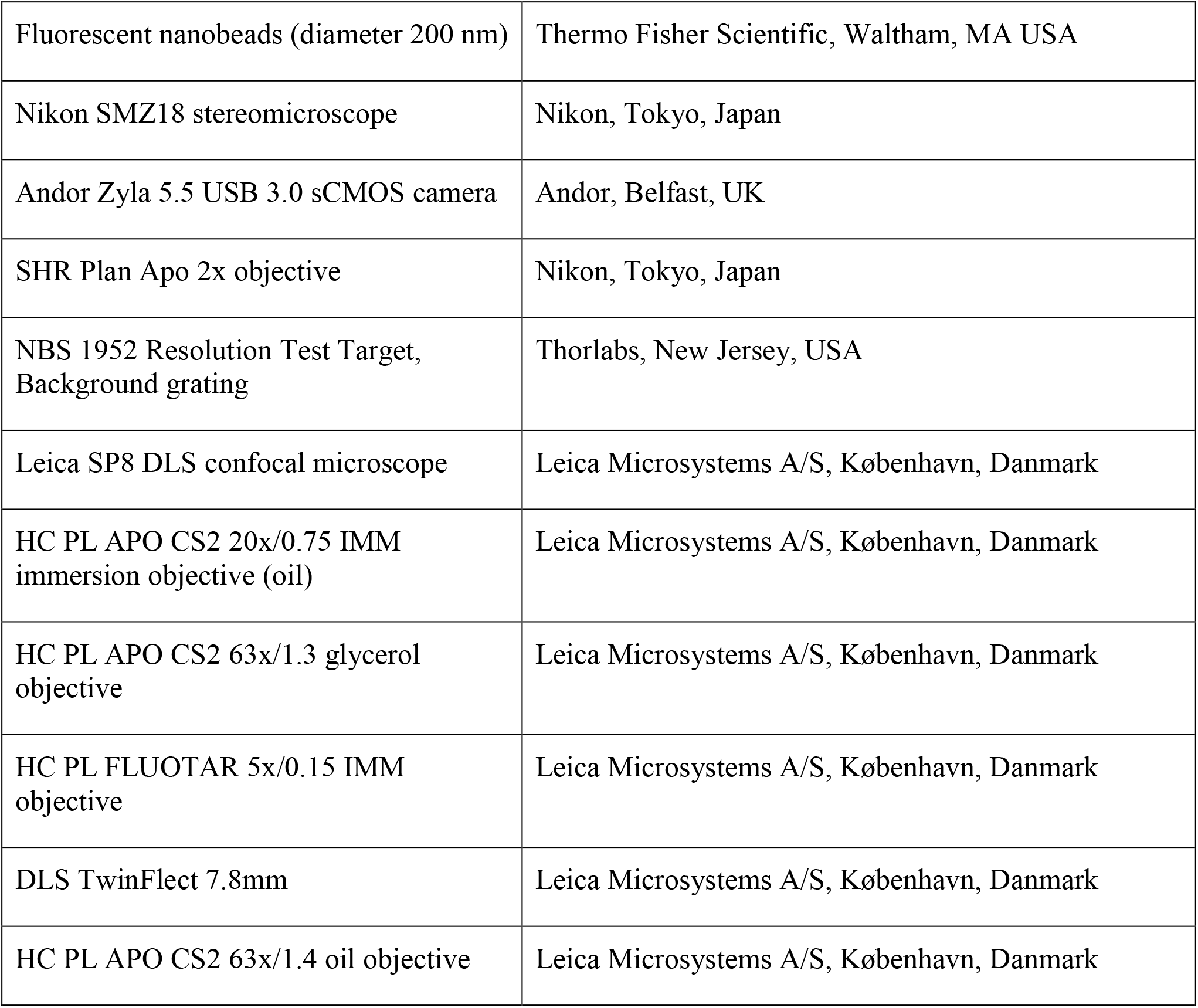

## 3 Methods

We developed clearing protocols to allow quantitative comparison of 2 variants of the iohexol based clearing medium SeeDB2 (abbreviated here as ‘SDB2G’, refractive index *n*=1.46, ‘SDB2S’ *n*=1.52); several variants of Rapiclear (hereafter referred to as ‘RC*x.x*’ where *x.x* is either *n*=1.47, 1.49 or 1.52), (c) 97% 2,2-thiodiethanol (‘TDE’, *n*=1.52) and (d) methyl salicylate in combination with permanent mounting in Permount (‘MS/P’, *n*=1.52). Where appropriate, we also tested the effect of pre-treatment to reduce air trapped in tracheal tubes, either by vacuum treatment or by dehydration through an ethanol series followed by rehydration.

### 3.1 Species used and dissection procedure

Animal species used were the large hoverflies *Vollucella pelluscens, Eristalis tenax* or the very similar *Eristalis pertinax*, and the blowfly *Calliphora vicina*. Although there is substantial variation between individuals, these all have similarly large brains (freshly dissected dimensions approximately 3000 by 1500 by 400μm, providing a suitable challenge for both objective lens working distance and the tissue clarity required for deep imaging. In total 71 flies were included in our analysis: 8 wild-caught *Volucella p.*, 23 wild *Eristalis sp.* (mostly *tenax*), 9 *Eristalis tenax* hatched from pupae purchased from Polyfly and 31 *Calliphora* hatched from pupae purchased from Blades Inc. Another single *Volucella p.* was used as an example application of intracellular dye injection for light sheet and high-resolution imaging. All animals/brains are labelled with an id (for example #01) which can be used to find meta-data in the supplementary table (see excel file S1 in supplementary data).

Many of the animals utilized were part of ongoing acute experiments and electrophysiological analysis or intracellular labelling of neurons for other projects, hence the sample of flies used are typical of real experimental scenarios and conditions. Following experiments, the animal was decapitated and the brain was extracted by dissection under 0.1 M phosphate buffered saline, pH 7.4 (PBS) and transferred into a small glass vial with 4% formaldehyde in PBS for fixation overnight at 4°C.

### 3.2 Pre-treatment

Following fixation, brains were first imaged to identify morphological landmarks and a pre-treatment estimate of tissue clarity (see 3.4 below). All images were captured using a Nikon SMZ18 microscope with an SHR Plan Apo 2x objective (working distance 20mm), with samples illuminated by diffuse incident light. Brains were then either pre-treated with ethanol (denoted by the subscript ‘E’ in figures and text below), vacuum (V), both (EV) or no pretreatment. Ethanol treatment consisted of dehydration and rehydration of the brain through an ethanol series diluted in PBS with steps of 50, 60, 70, 80, 90, 95, 100, 95, 90, 80, 70, 60, 50 % ethanol each for 20 minutes on a shaker at room temperature (RT). Note that the methyl salicylate /Permount clearing protocol (MS/P) always requires an ethanol dehydration series without subsequent rehydration since this medium is not miscible with water. In all other protocols, our application of the label E implies both dehydration and rehydration prior to clearing.

Vacuum treatment consisted of two 10-minute treatments with the brain immersed in the PBS in a 5ml glass vial with an open lid, placed in a vacuum chamber at a vacuum pressure of approximately 75 kPa. The PBS was changed between treatments.

### 3.3 Clearing and mounting

Following pre-treatment (if any) brains were then cleared according to the following protocols:

#### 3.3.1 Rapiclear (RCx.x)

After pre-treatment, brains were transferred into an eppendorf tube with 30-60 µL Rapiclear (RC1.52, 1.49 or 1.47) overnight on a shaker at RT. The brain was then mounted between two #1.5 coverslips (160μm thickness) using 500µm self-adhesive spacers (Sunjin labs) with fresh Rapiclear applied from a transfer pipette to fill the well created by the spacer. The coverslips were then attached temporarily to a glass microscope slide using lab tape to permit imaging from either side, if desired.

#### 3.3.2 Methyl salicylate & Permount (MS/P)

Following dehydration through an ethanol series, brains were transferred from the plastic vial or well plate to a 5ml glass vial, since methyl salicylate is a solvent for many plastics. A solution consisting of 50% ethanol and 50% methyl salicylate was added to the glass vial for 15 min. Finally, the brain was left on the shaker overnight at room temperature in 100% methyl salicylate. The following day brains were mounted between two coverslips using Permount. Plastic spacers (Avery Zweckform) were used rather than the self-adhesive spacers described above, since we found that the Permount reacted with the adhesive, generating additional bubbles in the mounted brain.

#### 3.3.3 SeeDB2 (SDB2G, SDB2S)

Following pretreatment, brains were cleared in a series of 33.3, 50 and 99.7% ‘Omnipaque’ solution with 0.3% Triton x-100 diluted in Tris-EDTA buffer. For each step, the brain was left on the shaker at room temperature overnight (or at least 6 hours for the first two steps). Omnipaque solution consisted of 56.2% iohexol (5-(N-2,3-Dihydroxypropylacetamido)-2,4,6-triiodo-N,N′-bis(2,3-dihydroxypropyl)isophthalamide, sold as ‘Histodenz’) and 43.8% Tris-EDTA. Tris-EDTA consisted of 10mM Tris-HCl (pH 7.6), 1mM EDTA and distilled water. The SDB2S protocol is the same as SDB2G but with a final immersion in 70.4% Histodenz solution instead of 56.2% (Ke et al., 2016). The brain was then mounted as for 3.3.1 above, using fresh Omnipaque solution.

#### 3.3.4 2,2-thiodiethanol (TDE)

The brains were treated through a dilution series (in PBS) series of TDE with increasing concentration of TDE by 10% every 30 min, to a final concentration of 97% (Staudt et al., 2007). The brain was then mounted as for 3.3.1 above, but using fresh TDE.

### 3.4 Transparency measurement and calculation

Images of each brain were captured before and after clearing using a Nikon SMZ18 stereomicroscope equipped with an Andor Zyla 5.5 sCMOS camera, using a 2x objective (working distance 20mm). Images were captured from two focal planes. The first was adjusted to allow dimension measurements from the brain itself (**Figure 1A,B**). The second image was with the focal plane adjusted to the plane of a background grating placed below the sample as it was viewed through the imaged brain (**Figure 1C**). Using ImageJ, adjacent rectangular regions of interest (ROIs) were selected within light vs dark parts of the images of the grating where it extended beyond the brain or where it was viewed through the imaged brain (**Figure 1C**). The average pixel intensity within these adjacent ROIs was used to estimate the average Michelson contrast in the image, *C*, where

**Figure 1.**
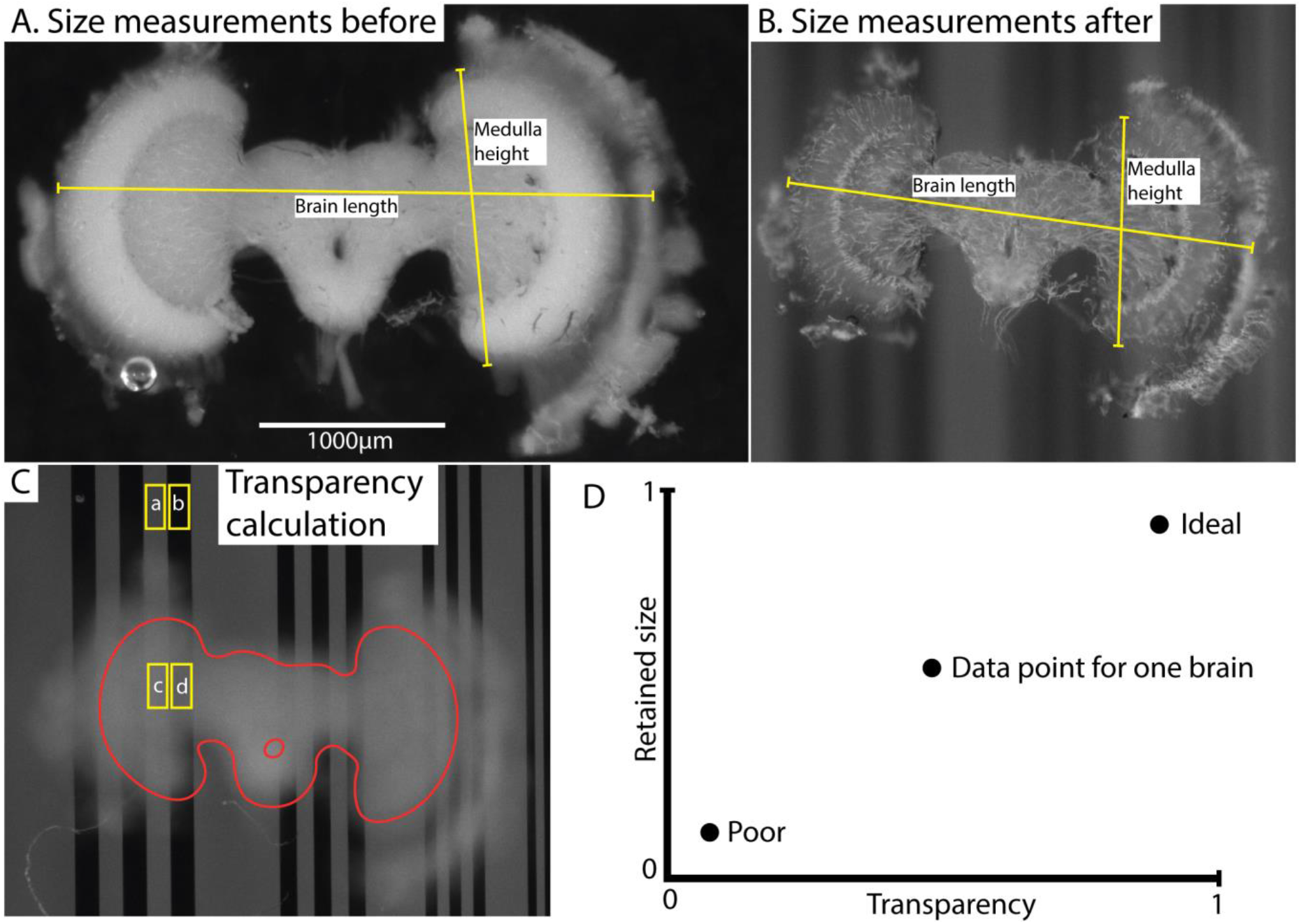
Illustration of measurement method (using brain #45) and results. **(A,B)** illustrates size measurements for brain length and medulla height before **(A)** and after **(B)** clearing. Although the measurements are based on medulla and brain height and length, the main principle was to use characteristic landmarks rather than actual anatomical structure length. The purpose of this principle was to enable length and height measurements on highly transparent brains. “Brain length” indicates the typical length used for comparison of shrinkage. “Medulla height” indicates the length used for normalization of transparency values. **(C)** shows selected areas (ROIs) used to calculate contrast; 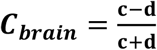 and 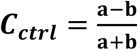, and transparency 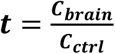. The labels (a, b, c, d) indicate the average grayscale value of the area. The brain structure (excluding lamina) is outlined in red. **(D)** illustrates the interpretation of the comparative data plot (which is used in **Figure 4A**). Transparency and retained size are the two dimensions, and ideal versus poor performance are indicated with two illustrative data points.

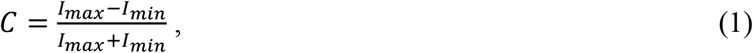

*I*_*max*_ is the maximum intensity (light) and *I*_*min*_ is the minimum intensity (dark). The transparency (*t*) was calculated with the following equation:

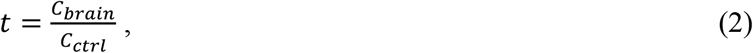

where *C*_*brain*_ is the contrast of the grating within the brain and *C*_*ctrl*_ is the contrast outside the brain.

Hence *t* is a measure of the contrast attenuation through the total thickness of the sample. To account for the varying brain size (and thus thickness in the Z dimension) between individuals, each transparency measurement (*t*) was normalized using the linear extent (*h*) of the medulla along the Y axis of the freshly fixed brain, a landmark that was consistently visible in every brain before clearing (as illustrated in **Figure 1A)**. The following formula was then used:

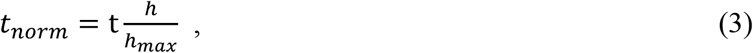

where *h* is the medulla height and *h*_*max*_ is the maximum medulla height measured among all brains analysed.

### 3.5 Length measurements for shrinkage estimation

The features used for linear dimension measurements were selected so that the measurement coordinates corresponded to the same part of the brain as well as possible in every animal. Due to the very high transparency in certain brains after clearing this was challenging at times. When possible, we measured the full width of the brain from edge to edge. **Figure 1A** shows “Brain length” which was typically used (the distance between the extent of the lamina/outer medulla in the left and right hemispheres) and “Medulla height”. Sometimes however, we were forced to use the only other landmarks clearly visible after clearing as reference points to measure the change in size, such as features of the margin of the dissected tissue or even individual tracheal tubes. Regardless of the landmarks used, we then computed the retained size (R):

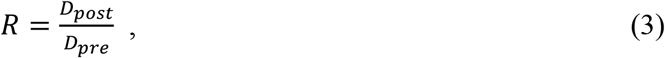

Where *D*_*post*_ is the length measurement obtained after pre-treatment and clearing, and *D*_*pre*_ is the length measurement obtained before treatment and clearing.

### 3.6 Additional experiments and example treatments

As part of our ongoing experiments, we intracellularly labelled several neurons (e.g. **Figures 5-6**) with a Lucifer yellow filled glass micropipette by injecting negative current (for details of the method, please see Barnett et al., 2007). The brains were then dissected, and fixed as above, then washed 4×10min in PBS, 55min in pure dimethyl sulfoxide, 3×30min in PBS with 0.3% triton X-100 (PBT), 3 hours with 5% natural goat serum (NGS) in PBT and then incubated for 3 days in biotin-conjugated anti-lucifer antibody diluted 50X in 2% NGS in PBT. The brains were then washed 3×30min with 10% NGS in PBT, and then incubated for 3 days in streptavidin-Cy3 diluted 50X in 1% NGS in PBT. Finally, the brains were washed 3×30min with PBS and cleared as described in section 3.3.1 above (e.g. RC1.49E for brain in **Figures 5-6**). Cleared brains were subsequently imaged using a Leica SP8 DLS confocal microscope and varying objectives and imaging modes. The microscopy images in **Figures 5-6** were obtained using this microscope in its confocal mode except **Figure 5C** that used the digital light sheet imaging mode (DLS, illustrated in **Figure 5B**). Confocal objectives used were either a 20x/0.75 (magnification/numerical aperture) immersion objective (using oil) or a 63x/1.3 glycerol objective. **Figure 5C** (DLS mode) was imaged using a 5x/0.15 objective and DLS TwinFlect 7.8mm mirrors both matched to the refractive index of glycerol (1.472).

**Figure 2.**
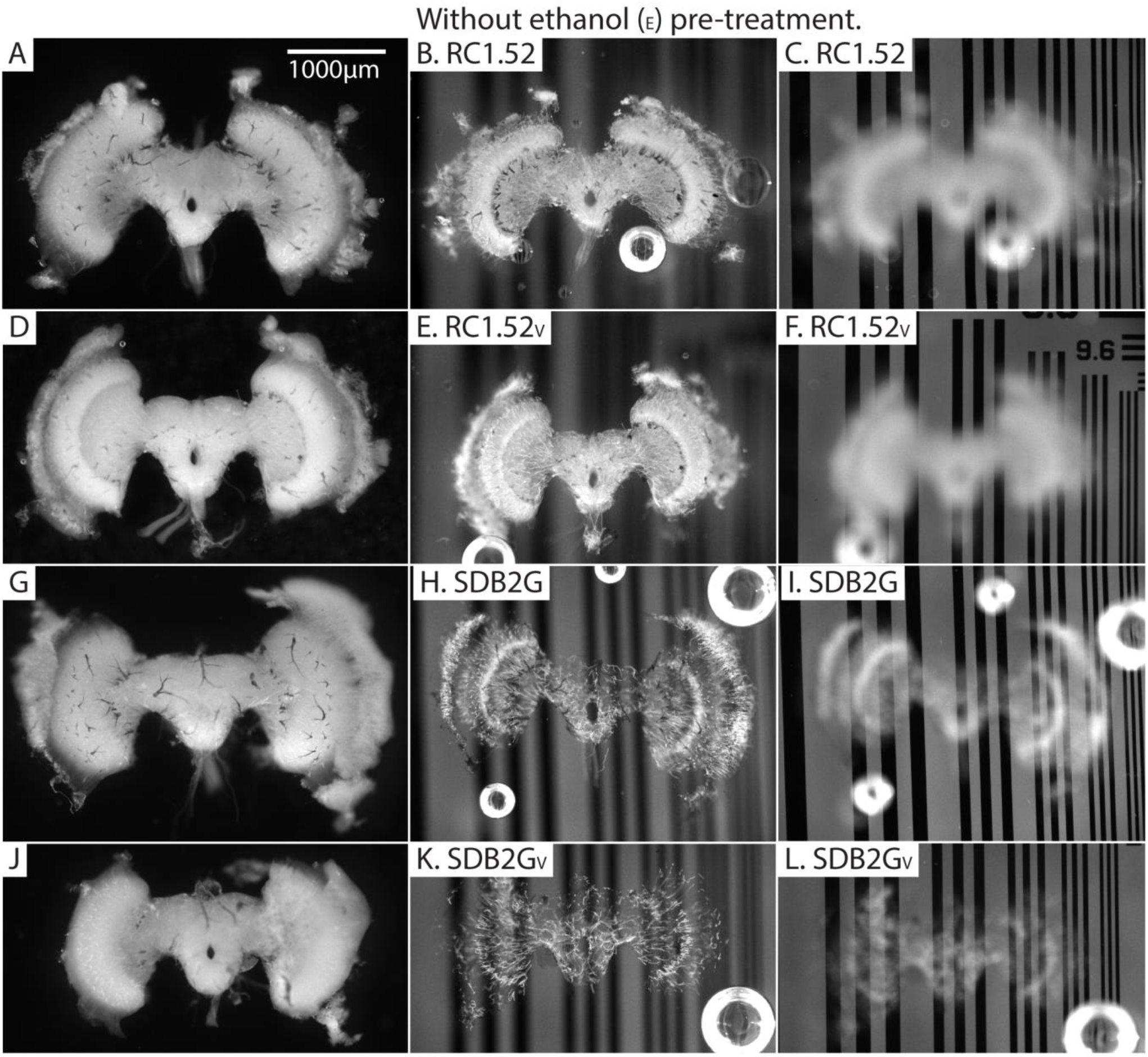
Pre and post clearing pictures of brains cleared with and without vacuum treatment (V). The uncleared brains are clearly visible in the left column and the cleared brains in the middle (brain in focus) and right (grating in focus) column are also visible. **(A-C)** is brain #56, **(D-F)** is #46, **(G-I)** is #67, **(J-L)** is #69.

**Figure 3.**
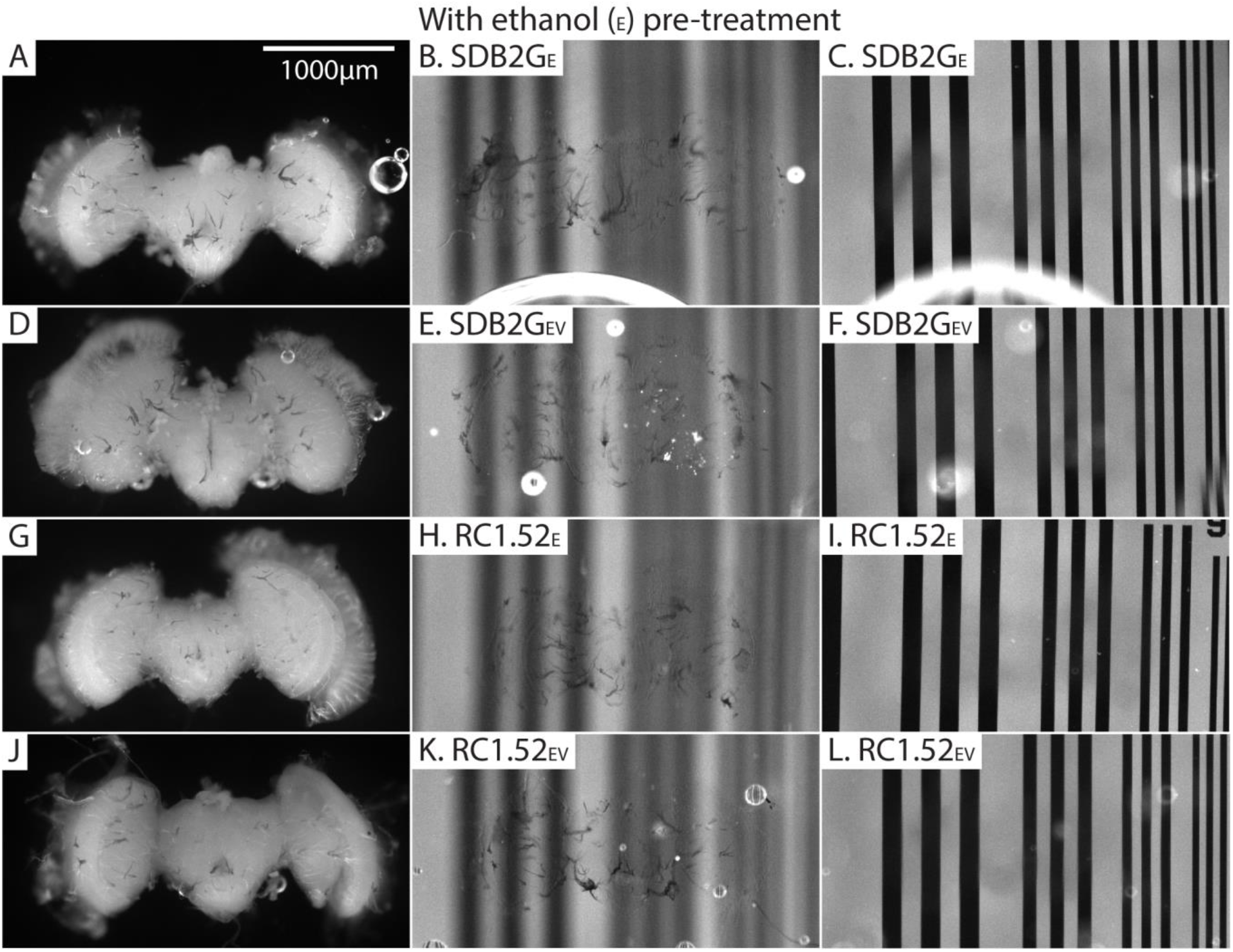
Pre and post clearing pictures of brains cleared with and without vacuum treatment (V) (all with ethanol treatment [E]). The uncleared brains are clearly visible in the left column and the cleared brains are barely visible (tracheal tubes visible) in the middle (brain in focus) and right (grating in focus) column. **(A-C)** is brain #80, **(D-F)** is #75, **(G-I)** is #82, **(J-L)** is #81.

**Figure 4.**
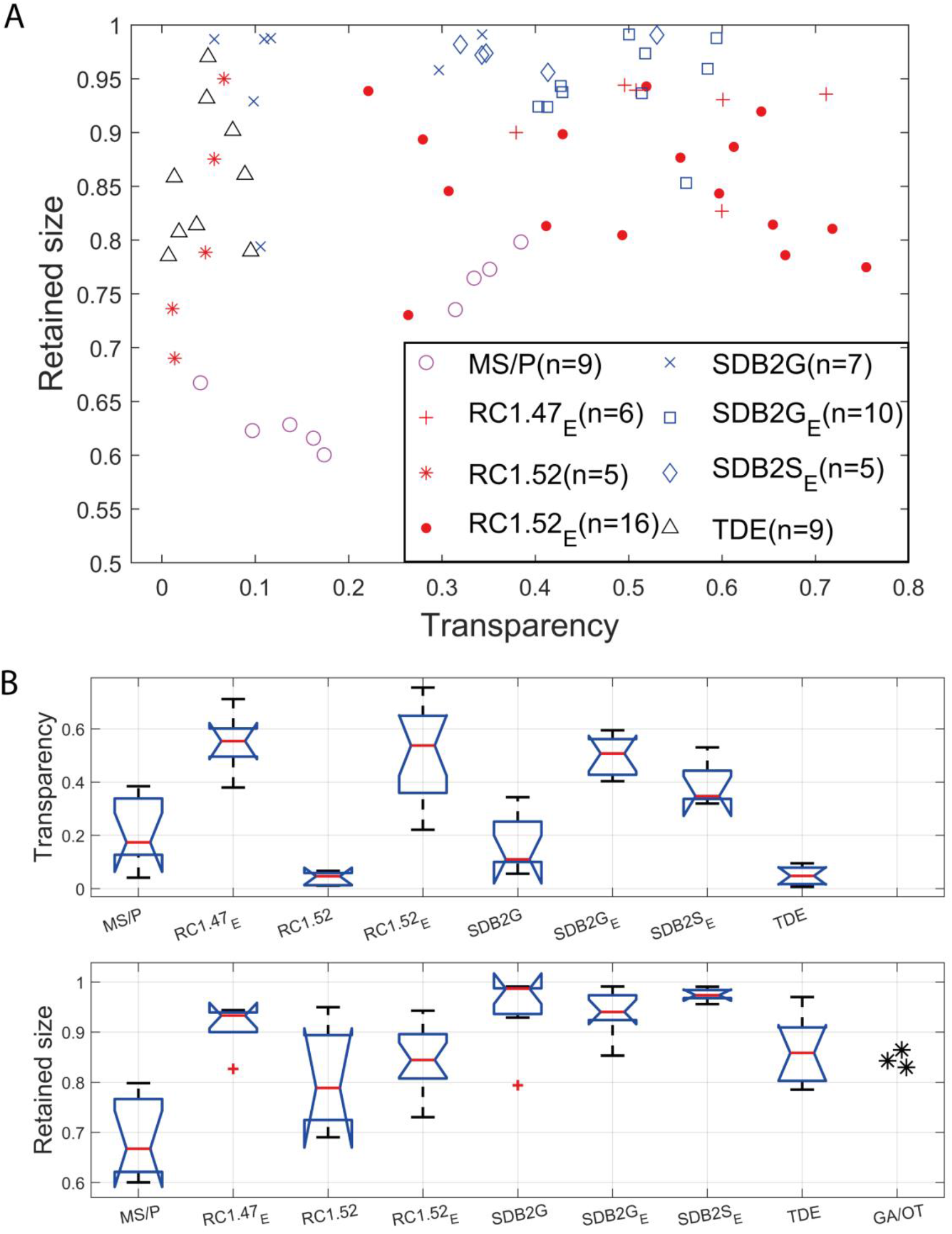
Clearing performance measured using transparency and retained size. **(A)** A scatter plot of the transparency versus retained brain size. (E) stands for ethanol treatment. The parenthesis after the clearing method name (n=…) indicates the number of samples (brains). **(B)** Box plot (Matlab R2016b built-in boxplot function) showing median (red line), notches, 25^th^ and 75^th^ percentiles (edge of boxes), whiskers (extreme data points). MS/P always includes ethanol dehydration but no rehydration. Each transparency measurement was normalized using the medulla height to compensate for differences in brain thickness.

**Figure 5.**
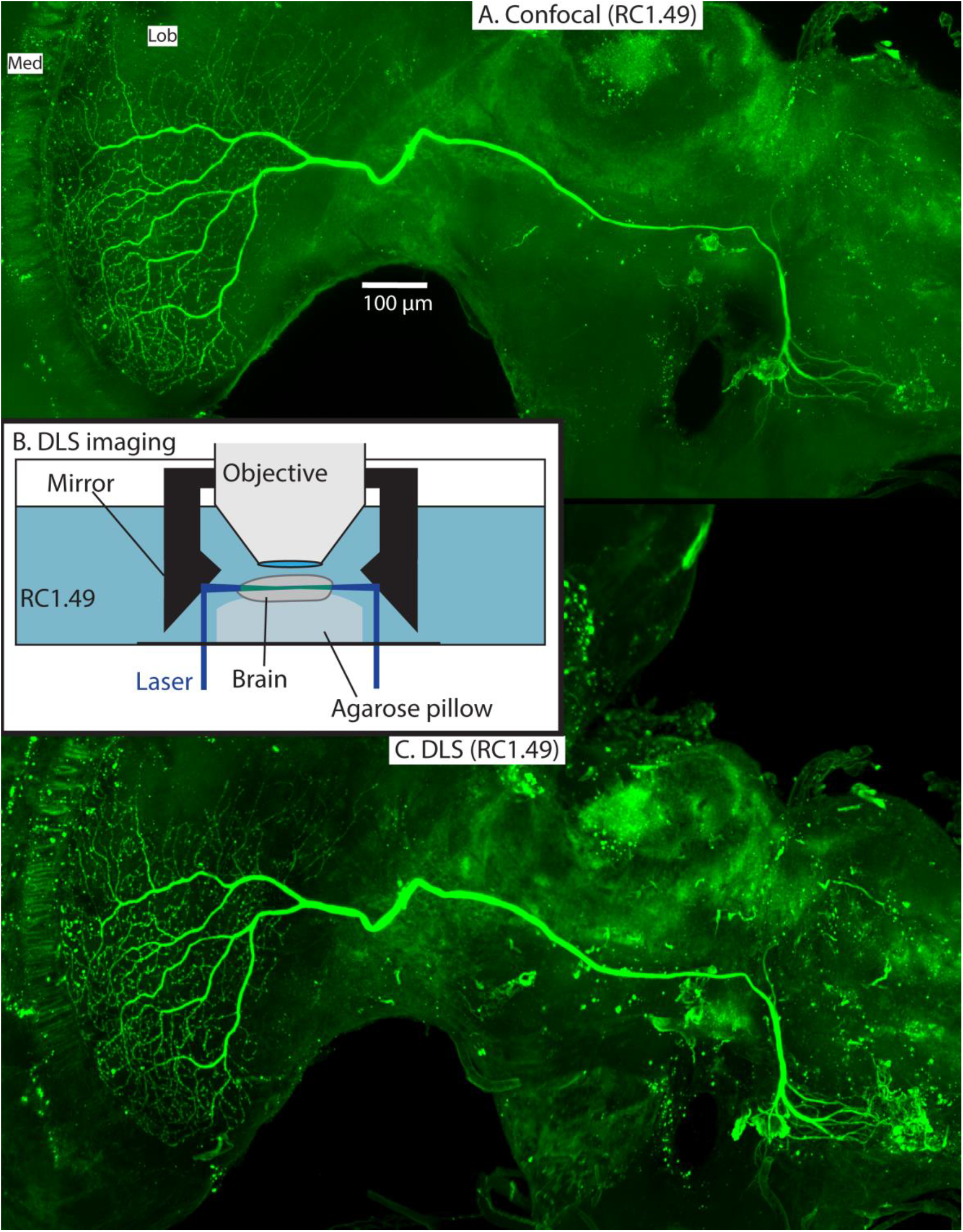
Example of a Lucifer yellow injected wide-field motion neuron in a brain (#160) cleared with RC1.49E. The neuron is a wide-field motion detector lobula complex neuron. The images were taken using a Leica SP8 DLS in confocal mode with a 20x oil immersion objective **(A)** and DLS mode with a 5x/0.15 objective and DLS TwinFlect 7.8mm mirrors **(C). (B)** illustrates the DLS imaging method used to acquire **(C)**.

**Figure 6.**
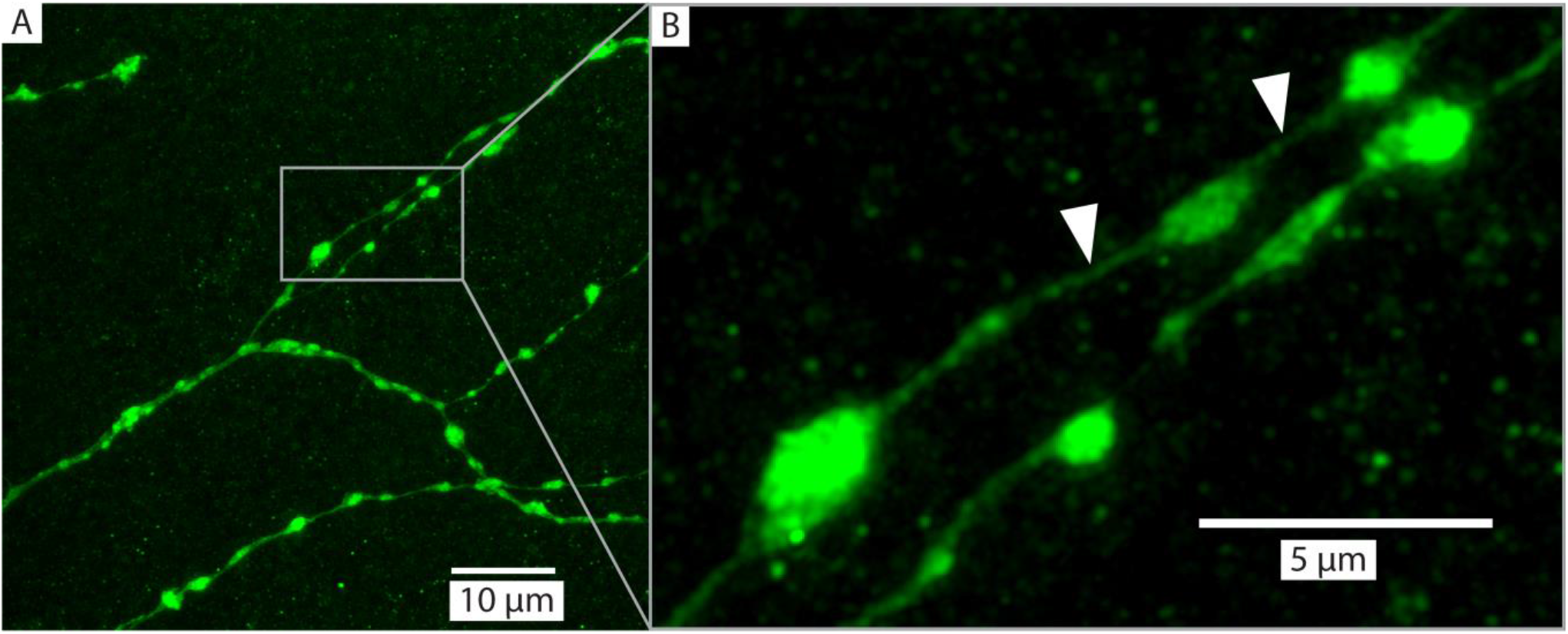
Example of a detailed branches of a tracer injected wide-field motion neuron [same as **(Figure 5)**] in brain (#160) cleared with RC1.49E and imaged with a glycerol objective (63x, NA: 1.3). The images were captured from a sample following 9 months of storage in room temperature. The imaging depth was around 50 μm bellow brain surface. **(A)** shows an overview of a group of branches and **(B)** zooms in on a subset to illustrate the details of a few ‘blebs’, which are considered to be an indication that the branches have output synapses (Hausen, 1976; O’Carroll et al., 1992). The white arrows indicate very fine neurites that have a diameter of between 136-271 nm.

### 3.7 Electron microscopy preparation protocol

An additional 3 *Eristalis tenax* brains were subjected to careful fixation as used in preparing samples for electron microscopy. Dissected brains were immersed in freshly prepared fixative solution, consisting of 2% paraformaldehyde and 2.5% glutaraldehyde in 0.1 M Na-cacodylate buffer (pH 7.4), for 24 hours at 4°C. The brains were post-fixed in 1% osmium tetroxide in distilled water for 2 hours at 4°C. The brains were dehydrated in graded ethanol series consisting of 70% ethanol (2×10 min), 96% ethanol (2×10 min), 100% ethanol (2×15 min). Brains were then transferred to acetone (2×20 min) as an intermedium before embedding first in dilute epoxy (Agar 100), with an acetone:epoxy mixture of (2:1) for 30 min, followed by 1:1 acetone:epoxy overnight, then infiltration with 100% epoxy for 8 hours before polymerization for 48 hours at 60°C. The final pictures were then photographed brain lengths measured as for the cleared brains.

### 3.8 Point spread function in Rapiclear using nanobeads

To estimate the 3-dimensional point spread function at different depths within Rapiclear, we prepared samples incorporating fluorescent nanobeads (diameter 200 nm). These were dispersed into low-melting agarose (1% in distilled water at 45°C) and then the warm solution mounted in a well formed using a spacer as for whole mount imaging (see 3.3.1 above) before clearing directly in the well using RC1.47, RC1.49, RC1.52. Confocal images (Z series) were then captured using a Leica SP8 DLS with a 63x/1.3 glycerol objective (RC1.47 and 1.49) and a 63x/1.4 oil objective (RC1.52).

## 4 Results

### 4.1 Qualitative clearing performance

Our protocol provides measurements of the effect of treatment (or pre-treatment) on changes in both brain dimensions (i.e. shrinkage or expansion) and transparency. Visual inspection of example brains cleared with RC1.52 and SDB2G but without ethanol treatment **(Figure 2)** clearly show that although more transparent than a fresh brain, the brains remain very visible as a cloudy foreground object, despite focusing on the background (**Figure 2 right column**). Vacuum treatment to evacuate air from the tracheal system (RC1.52V and SDB2GV) provided no obvious improvement in transparency over untreated brains (RC1.52 and SDB2G). However, pre-treatment by ethanol dehydration/rehydration (RC1.52E and SDB2GE) dramatically improved transparency (**Figure 3**) to the extent that brains cleared using these media are barely visible at all when the background is in focus (**Figure 3 right column**). Again, vacuum treatment made no apparent change to the transparency of ethanol pre-treated brains (RC1.52EV and SDB2GEV) suggesting that ethanol dehydration is a more effective way to displace air from these tubes that contribute to scattered light. Although it is hard to measure, vacuum treatment could be decreasing the thickness of the larger tracheal tubes, which may not affect overall transparency much (as the results indicate), but may still improve imaging locally. Visual examination of a few samples of SDB2G, SDB2GV, SDB2GE, SDB2GEV, RC1.52, RC1.52V, RC1.52E and RC1.52EV indicates no difference in reduction in the thickness of the tracheal tubes (**Figure 2 and 3**, black patches & lines). Note that the vacuum chamber we had access to was not capable of providing a powerful vacuum, and the protocol provided by the manufacturer of Rapiclear recommended repeating 10 minutes of vacuum treatment 4 times for insect tissue, while we only performed 2 cycles. Hence it remains possible that additional vacuum treatment may further improve evacuation of the tracheal system.

Finally, we note that there is no apparent difference in transparency between Rapiclear and SDB2G cleared brains following ethanol pre-treatment, with both achieving very high transparency (**Figure 3** right column). The qualitative results we obtained for Rapiclear with ethanol pretreatment are impressive considering the simplicity and speed of the clearing protocol (direct transfer from buffer to the clearing medium).

### 4.2 Quantitative clearing performance

**Figure 4A** shows results obtained after applying the clearing method to each brain and quantifying both the transparency using a metric that takes individual variations in brain size (and thus depth) into account, plotted against the retained size (as a relative fraction of the original size, 1.0) to account for shrinkage. In this representation, an ideal clearing medium would produce both perfect transparency and minimal shrinkage, leading to points lying in the upper right of the graph (see **Figure 1D**). The traditional protocols MS/P, TDE and the newer methods (RC1.47 and RC1.52) without ethanol treatment tended gave lower median transparency than RC1.47E, RC1.52E, SDB2GE and SDB2SE (summarized as Box and whisker plots, **Figure 4B**). Some clearing protocols, including MS/P and RC1.52E showed substantial variability in transparency between individuals (**Figure 4A**).

Although it gave little tissue shrinkage, TDE gave very low transparency compared to other the other clearing protocols, which surprised us due to the reported usefulness of this clearing medium (Gonzalez-Bellido and Wardill, 2012). In terms of the point-spread function (PSF) of nanobeads suspended in the medium, this was the best of the clearing media tested by Ke et al. (2016), although they did not report on absolute tissue clarity. Considering the reports that this medium also quenches fluorophores more than other protocols, it is difficult for us to recommend it for insect tissue, although perhaps tuning of the TDE concentrations or inclusion of ethanol pre-treatment could improve the transparency. That said, further addition of protocol stages would also make this protocol relatively complex compared with Rapiclear or SeeDB2 clearing, since it already involves progression through a slow TDE series. This could perhaps be amended if either the TDE and/or ethanol series were reduced by adding larger concentration increments at each step and reducing the waiting time between the steps, but given the superiority of the other techniques, we did not attempt further fine tuning of this protocol.

### 4.3 Refractive index dependent transparency and shrinkage comparison

Clearing of tissue using the 4 agents we tested here works mainly by matching the refractive index of the medium to that of the tissue in order to reduce scattering. Water soluble media such as SeeDB2 and Rapiclear permit refractive index tuning by varying the concentration of the non-aqueous components of the medium. We might, therefore, expect not only differences in transparency between different variants of the same medium, but also in shrinkage due to potential water extraction from the tissue at high concentrations of the medium.

For our insect brains, both ethanol-treated variants of SeeDB2 gave good transparency, but the higher index variant, SDB2SE, was actually worse than SDB2GE on average. The transparency performance for both RC1.52E and RC1.47E was among the best of the media tested and very similar to SDB2GE, although once again if anything the lower refractive index variant actually gave slightly better transparency (**Figure 4B**). While these slight differences may be due to partial mismatch between the medium and the average refractive index of the fixed brain, we should note that SDB2S in particular is a very viscous medium, more so than either Rapiclear variant tested or the less concentrated iohexol solution (SDB2G): Despite our application of a dilution series to aid penetration before immersing brains in the final concentration, it may be that SDB2S is less likely to infiltrate key structures such as small tracheal tubes than the less viscous media.

Shrinkage was not a serious issue for any variant of the iohexol based media (SDB2), with the vast majority of brains retaining greater than 90% of their original size (**Figure 4B**). Shrinkage of brains immersed in RC1.52E was around double that seen in the lower index version (RC1.47E) which, like the SDB2 brains, retained almost 95% of their original size. While the exact composition of Rapiclear is currently proprietary, it includes glycerol with a maximum refractive of 1.473 at 100% concentration. Hence it is possible that the very high concentration of other reagents required to achieve a higher index (*n*=1.52), leads to some water extraction from tissue and thus worse shrinkage compared to the lower refractive index variants. In terms of shrinkage, the widely applied MS/P technique was the worst protocol by far among those tested, with a median shrinkage to only 66.74% of the original size (**Figure 4B**). This gross change is immediately evident from the individual brain illustrated in **Figure 1A,B** before and after MS/P which we selected to illustrate our typical measurement landmarks.

It is likely that MS/P gives high shrinkage due to the ethanol dehydration (without rehydration) required before embedding, as well as through evaporation of the mounting medium solvent (toluene) following final mounting, which can also introduce unwanted bubbles and cavities into the space surrounding the brain. We wondered, however, whether this might also result in part from the limited crosslinking achieved by formaldehyde-only fixation. Most tissue fixation protocols for wholemount neuron imaging use relatively simple formaldehyde fixation in PBS, often using formaldehyde solutions made up from liquid stocks containing preservatives such as methanol to extend the shelf life, rather than freshly mixed paraformaldehyde (PFA) solution. While broadly compatible with many post-fixation treatments, such as immunohistochemistry, this may not provide optimal fixation and cross-linking of proteins and thus control over tissue shrinkage during subsequent dehydration. To provide an estimate of the minimum shrinkage possible if the same type of brain tissue is subjected to ‘gold standard’ optimal fixation, we therefore subjected a small number of brains to an electron microscopy (EM) preparation protocol. EM fixation combines both freshly mixed paraformaldehyde with the more strongly protein crosslinking fixative glutaraldehyde (GA), as well as post-fixation with osmium tetroxide (OT). The latter preserves lipid structures which would otherwise dissolved during alcohol dehydration following formaldehyde only fixation, potentially contributing to shrinkage during subsequent dehydration/embedding. Based on 3 GA/OT fixed and epoxy resin embedded brain samples we prepared, the average shrinkage was indeed substantially less than in MS/P mounted brains, with a mean retained size close 85% (see asterisks in **Figure 4B**, lower). It would require further work to establish whether this improvement following dehydration is due only to the fixatives used, or also to the non-solvent based embedding medium (since epoxy hardens though polymerization rather than solvent evaporation), or both. Nevertheless, while this gave an improvement over MS/P embedding, the ‘gold-standard’ fixation treatment still clearly results in more shrinkage than we obtained with the best of water-miscible clearing media such as SeeDB2 or Rapiclear (**Figure 4B**).

### 4.4 Compatibility with large scale and super-resolution imaging techniques

Our analysis above shows that some of the more recently introduced clearing protocols with a lower refractive index (∼1.47), that better matches insect brain tissue, have great potential for high quality whole-mount imaging. Rapiclear in particular involves a very simple sample preparation protocol (direct transfer to the medium from buffer following fixation and pre-treatment). In order to demonstrate the usefulness of this technique in real-world applications, we cleared a number of brains from the hoverfly *Eristalis tenax* in Rapiclear following electrophysiological recording and intracellular injection of single neurons in the optic lobes with the fluorescent probe Lucifer yellow (see 3.6). We identified a number of large ‘tangential’ neurons in these brains with neural processes ramifying throughout a range of depths within the outer lobula complex and then with projections of the axon 1mm or more across the brain to contralateral midbrain structures. The large size of such neurons and the range of depths of their processes makes them challenging to image at high resolution in whole mount.

**Figure 5** illustrates an example from one of the stained brains. This was a wide-field motion sensitive neuron with high sensitivity to downwards and rightward moving bars and grating patterns in the contralateral side of the visual field, suggesting that its inputs were the dendrites visible in the contralateral midbrain. The z-projection of the neuron in **Figure 5A** was obtained in the confocal microscope within a day of commencing the clearing process, illustrating the speed of the technique and the high quality of resulting images. In this case we used an intermediate refractive index (RC1.49) to provide a compromise between the slightly greater shrinkage of RC1.52 and the ideal medium for a 20x oil immersion objective, which gave us enough working distance to image through the full depth of the brain.

**Figures 5C & 6** show the same sample as **Figure 5A** after 9 months of storage at room temperature (in darkness). For **Figure 5C** we exploited an additional advantage of the non-hardening medium by first removing one coverslip and simply rinsing off the old Rapiclear (which remained fluid) before re-mounting the sample so that it was suspended on an agarose pillow above the base of a glass-bottomed imaging well (**Figure 5B**) and then re-clearing overnight in fresh RC1.49. We then used the Leica SP8 DLS microscope in the ‘Digital Light Sheet’ (DLS) mode, which reflects a scanned light sheet into the brain from the sides using a pair of mirrors attached to the imaging objective, which in this case was immersed directly in the Rapiclear. Both the objective and mirror set were optimized for working in clearing media with the refractive index of Glycerol. For direct comparison with the confocal series in **Figure 5A**, we still imaged the brain in the same orientation as the confocal series, i.e. with the objective directed at the posterior brain surface and the Z axis corresponding to the anterior-posterior axis of the brain (‘Posterior view’), which is the thinnest overall dimension of the insect brain. This imaging mode places high dependency on the degree of clearing, since excitation laser light has to pass through larger horizontal dimension of the sample e.g. through the ventral and dorsal margins of the medulla in order to excite fluorescence within the lobula. Any residual opacity in the medulla (or uncleared tracheae) casts shadow artifacts onto the resulting horizontally illuminated columns as the laser scans the light sheet. Despite the absolute resolution limits of the lower N.A. objective used for the DLS images, the overall quality of the image stacks was still excellent compared with the confocal images, especially considering the total imaging time required was only a few seconds (**Figure 5C** and **supplementary video S2).**

### 4.5 Super-resolution imaging at large depths in Rapiclear

Upon the introduction of SeeDB2S and SeeDB2G, the authors who developed these media already highlighted their potential for application of a number of super-resolution imaging techniques. To explore the similar potential for Rapiclear, we obtained super-resolution images (**Figure 6)** from a detailed region of the lobula plate arborization of the same sample as in Figure 5A, again re-imaged 9 months after the initial clearing. These were obtained using the Leica SP8 microscope in “Hyvolution” mode, with a 0.5 Airy unit pinhole and 2x oversampling in X, Y and Z, and a high numerical aperture 63x objective matched to glycerol immersion. Following acquisition, the image stack was first stabilized for 3-dimensional drift, before application of deconvolution using the high-resolution options in the Huygens software that provides the core to the Hyvolution mode. These images clearly resolve many very small diameter processes within the terminal dendrites (e.g. **Figure 6B** white arrows, neurite diameter 136-271nm) lying between varicosities (‘blebs’) that are most likely clumps of mitochondria, as typically associated with output synapses in insect neurites (O’Carroll et al., 1992).

The excellent image resolution evident from **Figure 6** shows that the transparency and the optical match between the medium and objective must, in practice, be sufficient to achieve a high-quality point spread function (PSF), even at substantial depths within the sample. Since this has not been quantified previously for Rapiclear, we further estimated the effect of image depth on the PSF by imaging fluorescent nanobead samples prepared by first suspending in 1% low melting agarose and then cleared and mounted as for the insect brains (using either RC 1.47, 1.49 and 1.52). These were imaged either using a 63x objective for glycerol immersion (for RC 1.47, 1.49, objective NA=1.3) or a 63x oil immersion objective (for RC 1.52, objective NA = 1.4) to try and optimize the match between clearing medium and objective immersion medium. The limited free working distance for the optimized oil immersion (*n*=1.52) condition limited us to imaging beads no deeper than 55µm below the coverslip. At this depth, the PSF is elongated in the Z dimension (**Figure 7**), but otherwise comparable to a bead 10µm below the coverslip. While minor, this elongation was somewhat surprising considering that the RC1.52 medium should be an index match for oil immersion. It is possible that this is due to small amount of residual water retained within the cleared agarose suspension in which nanobeads were embedded to restrict their Brownian motion.

**Figure 7.**
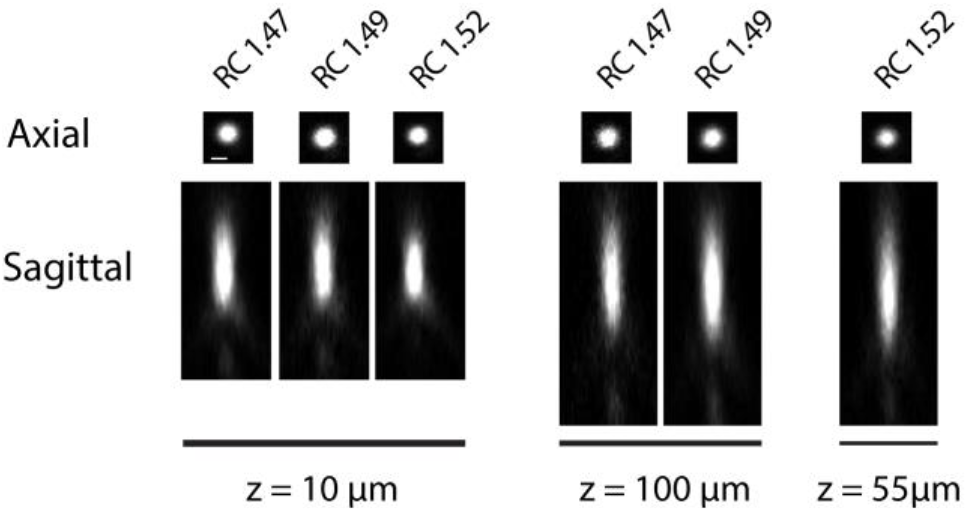
Point spread functions of fluorescent nanobeads (diameter: 200 nm) embedded in agarose and RC1.47, RC1.49 (63x, NA: 1.3) and RC1.52 (63x, NA: 1.4) with different refractive indices and at different sample depth (z).

At shallow depths (10µm) PSFs for the lower index Rapiclear variants are slightly more elongated in the Z dimension than in the oil immersion (1.52) condition, consistent with the higher numerical aperture of the latter objective. Importantly however, at 100 µm depth the PSFs for these media were similar to that obtained by oil immersion, with no obvious additional elongation along the Z axis, despite the additional (almost 2x) depth. Considering our earlier observation (**Figure 2B**) that the tissue clarity of both SeeDB2S and Rapiclear 1.52 were worse than their glycerol matched counterparts, it is likely that the effective refractive index of the brain tissue once fixed and cleared in these media is closer to 1.47 than to 1.52. Hence while the additional numerical aperture available for oil immersion may provide better resolution for very shallow samples, our results suggest that the best results for super-resolution imaging in very deep insect brain samples will come from applying a lower index clearing medium in combination with an objective matched to glycerol as the immersion medium, as we used in **Figure 6.**

## 5 Discussion

In this paper, we have limited our analysis to a small number of clearing media, including the recently developed SeeDB2. In introducing the latter as a good alternative for super-resolution applications, Ke et al. (2016) used Z stacked images of fluorescent nanobeads to compare the PSFs of a number of additional clearing media that have been widely employed across a range of species, particularly rodents. Their results confirmed a well-controlled expansion of the PSF at depths of 100 µm for both TDE and SeeDB2, which we also see here for Rapiclear (not included in their analysis). However, a number of other widely employed media included in their analysis gave clearly inferior PSFs by a depth of 100µm and correspondingly more rapid fall off in brightness with imaging depth. These included Scale/S, ProLongGold, FocusClear, and CUBIC. In our tissue, we found that Rapiclear gave similar clarity and shrinkage to SeeDB2, but is even easier and more rapid to apply.

Our results and protocols provide objective grounds for other researchers to select between the tested clearing media for the one best suited to their specific application. However overall, our results suggest that the two recently introduced clearing media, SeeDB2G and Rapiclear 1.47, provide a superior degree of tissue clarity, tissue shrinkage control and potential for high resolution imaging at large depths within insect brain samples, compared to traditional clearing media used by many labs. The reduced tissue shrinkage in particular should be a serious consideration for any high-resolution application. Assuming that neuron structures shrink isometrically, the large shrinkage we quantified for the traditional methyl salicylate/ Permount samples represents a very substantial loss of potential resolution during subsequent imaging, irrespective of the objective used. Although of course this is partially offset by having a smaller volume that requires imaging (and thus reduced objective working distance).

If long term storage or perfect mechanical stability during imaging are key considerations, there may still be compelling reasons to use permanent hardening mounting media such as Permount, for example. Although we have not yet had an opportunity to evaluate the very long-term storage potential of the Rapiclear technique, we have a number of samples stored simply in the dark exactly as initially prepared (i.e. mounted between two coverslips using a 500µm self-adhesive plastic spacer) for well over a year at room temperature, and which still show excellent fluorescence. Indeed, the image in **Figure 6** was obtained from the identical sample to **Figure 5A** after 9 months of storage at room temperature. Nevertheless, MS/P is known to be safe for storing samples for years with preserved fluorescence and tissue quality.

### 5.1 Variability in transparency and shrinkage

Some clearing protocols, including MS/P and RC1.52E showed substantial variability in transparency between individuals (**Figure 4A**). This may in part reflect differences between species included in the analyses, even though we normalized this measure to account for variability in brain size. Potentially, certain species could be more easily cleared with some clearing methods, due to differences in tissue structure (including tracheal distribution). Such variation might also explain the poor performance of TDE in our study using Dipteran brains compared with the good clearing reported for this medium in an earlier study on dragonfly CNS tissue (Gonzalez-Bellido and Wardill, 2012). These differences may also result in part from variations in the dissection procedure (carried out in this case by more than one researcher), where some brains were left with more superficial tracheal tubes and air sacks left on the brain, as can be seen in the uncleared brain in **Figure 2G**, for example (*cf* **Figure 2D, J**). This could affect transparency within the selected ROI since this was typically placed within the lobula, where such tracheae are prominent (see **Figure 1C**).

### 5.2 Deep tissue imaging techniques

In emerging deep-tissue imaging applications such as light-sheet imaging, a further advantage of aqueous miscible clearing media is their compatibility with a range of direct immersion objectives. The high degree of additional tissue transparency that we obtained with such media is another compelling reason to consider them for such applications. The tissue clarity was certainly adequate to get uniformly illuminated light sheet images of an adequate quality to allow for neuronal morphology 3D reconstruction for confirmation of neuronal identity or neuron subtype identification. Clearly the simplicity and speed of this clearing technique offers some potential advantages for high-throughput analysis of neuronal structure compared with more traditional images obtained from multiple Z series by confocal imaging (which may take many hours per sample to acquire). Indeed, with the Leica SP8 DLS microscope system we used in this study, it is even possible to achieve near simultaneous imaging of the whole brain from above in the DLS mode, and confocal imaging from below (at least using air or water immersion objectives with a working distance long enough to reach the sample through the agarose pillow). Of course, confocal images are also directly available by simply remounting the sample between coverslips (which takes minutes) if even higher resolution imaging of key structures is required using shorter working distance objectives e.g. for more detailed reconstructions for computational modeling studies where the interest lies in the fine neurites.

One downside to these new media is that the reagents involved in both are relatively expensive (either proprietary in the case of Rapiclear, or based on products made for human clinical use in the case of the Omnipaque used in SeeDB2), at least compared with methyl salicylate or TDE. This may be a consideration for light sheet microscopy in particular, where the objective is immersed in the same medium, requiring a relatively large volume. In our own hands, we have found that this is mitigated by re-cycling some of the immersion medium for future imaging sessions, as well as by other advantages such as the ease of rinsing excess Rapiclear from the objectives afterwards in water. SeeDB2 is less suitable in this regard, as it tends to harden rapidly on contact with air.

### 5.3 Toxicity

The preparation of samples using the newer media (both SeeDB2 and Rapiclear) is rapid and straightforward. SeeDB2 is not considered to be seriously toxic (Schega et al., 2020), although can have adverse effects when injected intravenously in large doses as an X-ray contrast agent in humans/animals (Stuart et al., 2009), thus favoring this protocol by contrast to MS/P, with less risk of exposure to highly noxious smelling or toxic solvents such as methyl salicylate, and the toluene in Permount, which must be handled at all times in a fume cupboard. The chemical compounds of Rapiclear are not fully known due to pending patent (SunJin Lab Co.). The MSDS for this product does note the presence of a small amount of preservative (Sodium azide) as a potential hazard. Although since this is a commonly used additive in many of the immunohistochemical procedures frequently carried out in insect brain research prior to clearing anyway, it requires no special precautions beyond those generally already employed in most labs. The fact that these clearing media have low toxicity and are safe to use on the open bench makes it easier to think in terms of robotic automation of experimental protocols which is becoming possible even for labs with lower sample output frequency (Savall et al., 2015; Gome et al., 2019; Poulsen et al., 2020).

### 5.4 Implications of a less viscous mounting medium

As noted earlier, SDB2G, RC and TDE are less viscous compared to Permount (particularly once the latter is fully hardened by solvent evaporation). This has pros and cons. Obviously a less viscous medium may be better for penetrating thick volumes as well as generally easier to handle during the mounting process. However, during the very lengthy (2x oversampling) imaging required for super-resolution techniques such as 3D deconvolution that we applied for **Figure 6** we noted a slight drift in the sample position over time. While such minor drifts are common due to small temperature changes leading to stage expansion / contraction, it is possible that the less viscous mounting medium for our sample contributed to this, especially since our microscope uses an inverted objective pathway, requiring the sample to be positioned upside down compared to its storage position. Although our software (via the Huygens image stabilization plugin) was able to correct for this drift it could be worth considering storing such samples in their imaging position a few hours before imaging to minimize this type of drift. Alternatively, the thickness of the spacer used to suspend the brain between the two coverslips could be reduced so that the brain is gently compressed along the Z dimension. On the plus side, one benefit of using less viscous mounting medium as demonstrated here is that it is easy to remove the sample and remount it in another orientation if required (**Figure 5B**).

## 7 Author contributions

Conceptualization: DOC, BB, HF. Clearing experiments and measurements: BB, HF. Tracer injection: BB. Confocal imaging: BB, DC. DLS imaging: DOC. Bead experiments: ER, DOC. Manuscript writing: BB, DOC. Comments on figures & writing: DOC, ER, HF.

## 8 Acknowledgements

We thank Seán McGovern for his help with brain dissections and data registration and Ola Gustafsson for help with the confocal microscope and electron microscopy sample preparation.

## 10 Data Availability Statement

Data is available upon request.

